# Vascular normalization in breast cancer confers an endothelial immune suppressive gene expression profile

**DOI:** 10.64898/2026.06.17.733057

**Authors:** Liqun He, Michael Welsh

**Affiliations:** Department of Immunology, Genetics and Pathology, Uppsala University; Department of Medical Cell Biology, Uppsala University

**Keywords:** Vascular leakage, vascular normalization, immune suppression, breast cancer, gene expression, *Shb*-gene

## Abstract

Tumors commonly exhibit a leaky vasculature and innate and/or adaptive immune cell infiltration. A recent study showed that a non-leaky and normalized mouse breast cancer vasculature after conditional deletion of the *Shb* gene in endothelial cells (EC) displayed a suppressive immune cell profile. To extend those findings further, regulation of immunomodulatory gene expression of normalized breast cancer EC was related to healthy human breast and human breast cancer EC gene expression. A considerable correlation was observed in the gene expression profiles of immunoregulatory genes between normal breast arterial EC and normalized (*Shb*-deficient) EC, and these changes did not primarily seem to involve gene products forming conduits for tumor leukocyte transendothelial migration but other aspects of immune suppression as for example cytokine expression. In addition to breast cancer, healthy lung and colonic EC exhibited an immune suppressive phenotype compared to their lung cancer and colorectal cancer counterparts. It is concluded that normalized EC may confer a non-leaky and immune suppressed phenotype, an insight that may be exploited to enhance the efficacy of immunotherapy.

## Introduction

Tumor immune responses are connected to vascular function (1). Particularly vascular normalization, *i*.*e*. the reduction of tumor vascular leakage and increase in vascular tumor perfusion commonly occurring in response to angiogenesis inhibition, has been shown to improve the efficacy of immune therapy in several different types of cancer (2, 3). Breast cancer has so far not been particularly amenable to immune therapy although triple Positive breast cancer may under certain conditions respond to such treatment (4).

A recent study on breast cancer explored the relationship between vascular function and immune response by conditionally deleting *Shb* in endothelial cells (5). The Src-homology 2 domain protein B (SHB) has been shown to be required for vascular endothelial growth factor-A (VEGFA)-induced vascular leakage and angiogenesis (6). An experimental model of E0771.lmb triple-Positive breast cancer in mice with endothelial *Shb*-deficiency was found to display decreased tumor vascular leakage, in line with vascular normalization, and tumor immune suppression (7). A human cohort confirmed a relative increase in regulatory T cell (Treg) numbers in tumors with less leakage (5). This observation was investigated further by performing bulk RNAseq on tumor endothelial cells (EC) and single cell (sc) RNAseq on tumor immune cells after conditional deletion of *Shb* in EC. The scRNAseq data revealed evidence for immune suppression and altered cytokine expression profiles whereas the endothelial gene expression changes detected by bulk RNAseq were in agreement with reduced vascular leakage and angiogenesis, features further supporting that a state of vascular normalization had been achieved (5). In that study, the analysis mostly focused on endothelial gene expression alterations pertaining to the formation of conduits allowing immune cell transendothelial migration (TEM). Several endothelial gene expression changes that influence tumor invasion of immunosuppressive immune cells were noted that potentially could explain the relative increase in Tregs (5). Our present data show a correlation between an immune suppressive phenotype of normalized mouse breast cancer EC gene expression and that of healthy breast arterial EC, suggesting a linkage between vascular normalization and gene expression changes that promote immune suppression.

## Methods

To expand the list of genes that may influence an immunosuppressive phenotype, the MGI-GO browser (https://www.informatics.jax.org/vocab/gene_ontology) was used to collect genes according to the biological processes “positive regulation of T cell activation”, “activation of immune response” and “cytokine activity”. The genes belonging to these categories were then used to extract genes from the list of genes with uncorrected p-values of less than 0.05 in response to *Shb* iECKO (conditional endothelial knockout). The extracted genes were then subjected to a second round of screening by gene ontology analysis at (https://davidbioinformatics.nih.gov/home.jsp) at which genes of relevance for immune related processes were collected (Table S1). Only genes expressed in EC that could exert an influence on immune cell function such as genes coding for proteins expressed at or outside the cell surface, cytokines and other secreted proteins were curated. Transcription factors at (https://www.informatics.jax.org/vocab/gene_ontology) were compared with changes of p<0.05 in the endothelial *Shb* iECKO/wild type and control/breast cancer EC gene expression arrays (Table S2). Transcription factor differences shared between the *Shb* iECKO and control breast EC comparisons were then used to find possible enhancer binding sites for these among the immune regulatory genes as identified by (https://www.genecards.org) in Table S2.

Human breast cancer, lung cancer and colorectal cancer scRNA-seq data were obtained from previously published studies (8-10). Annotated vascular EC populations were extracted for downstream analysis. Differential gene expression analyses between control and tumor samples were performed separately for arterial, capillary (type I and II for lung and colorectal cancer), and venous ECs, as well as for the pooled EC population, comparing control versus tumor conditions.

Human atherosclerotic plaque scRNA-seq data were obtained from a published study (11). Annotated EC populations were analyzed, including three EC clusters: ECs, EndoMT ECs, and pro-angiogenic ECs. Differential expression analyses were performed by comparing ECs against EndoMT ECs and pro-angiogenic ECs, respectively.

Differential gene expression analyses were conducted using the FindMarkers function in the Seurat R package (version 4.3.0). This function applies the Wilcoxon rank-sum test to assess differential expression, followed by multiple-testing correction using the Bonferroni method. Genes with an adjusted P value < 0.05 were considered significantly differentially expressed. To identify potential transcription factors regulating the differentially expressed genes, transcription factor binding site enrichment analysis was performed using the RcisTarget package (version 1.18.2) in R with default parameters. This approach identifies over-represented transcription factor binding motifs within the gene sets, which are subsequently annotated to their corresponding transcription factors.

Significances of correlations by Pearson’s R coefficients between groups were determined in Prism 11™. Mouse EC gene expression changes (uncorrected p-values to prioritize possible changes) of bulk RNAseq were taken from (5). In some comparisons, only relative changes (regardless of p-values) were used. For gene ontology (GO) analysis of gene expression differences between normal (healthy) and cancer human EC, corrected p-values less than 0.05 were used. GO p-values were corrected for false discovery rates by the Benjamini-Yekutieli procedure.

All data presented on comparisons between *Shb* iECKO EC and normal human EC were calculated as *Shb* iECKO over wild type EC ratios versus normal (healthy) human EC over breast cancer EC ratios for arterial, capillary and venous populations, respectively, and denoted in the text as *Shb* iECKO relative normal human EC gene expression.

## Results

In order to obtain a better understanding of the immunoregulatory status of normalized breast cancer endothelium, we conducted an additional search for immunomodulatory genes in our previously published data-set as described in the methods section and identified 10 more genes besides those already described (5). These and some of the previously noted genes (5) are presented in Table S1 with changes in expression indicated by red (increase) and blue (decrease) and were subjected to a literature search (Table 1). Based on literature, *Cd5l, Cd36, C2* and *Vegfa* appear to primarily affect inflammation and innate immunity in the short term (12-15). The reduced *Shb* iECKO EC gene expression changes of these genes will probably cause a less inflammatory and thus suppressed immune environment. Long-term exposure to VEGFA appears to inhibit dendritic and effector T cells (16). The gene expression changes of *Bmp2, Lgals9, C1qbp, IL24, Icosl* and *Msmp* are all predicted to be primarily immune suppressive, either by exerting effects on T effector or Treg cells (17-24). Complement factor H (*Cfh*) inhibits T effector cells (25) and its reduced expression is predicted to be immune stimulatory which together with the long-term effects of VEGFA are the only changes suggesting a predominantly immune stimulatory response. In summary, the currently curated set of genes provides additional support for EC promoting immune suppression upon *Shb* gene deletion in addition to the previous study demonstrating a relative increase of tumor Tregs and examining endothelial gene expression changes primarily related to immune cell TEM (5).

**Table 1:**
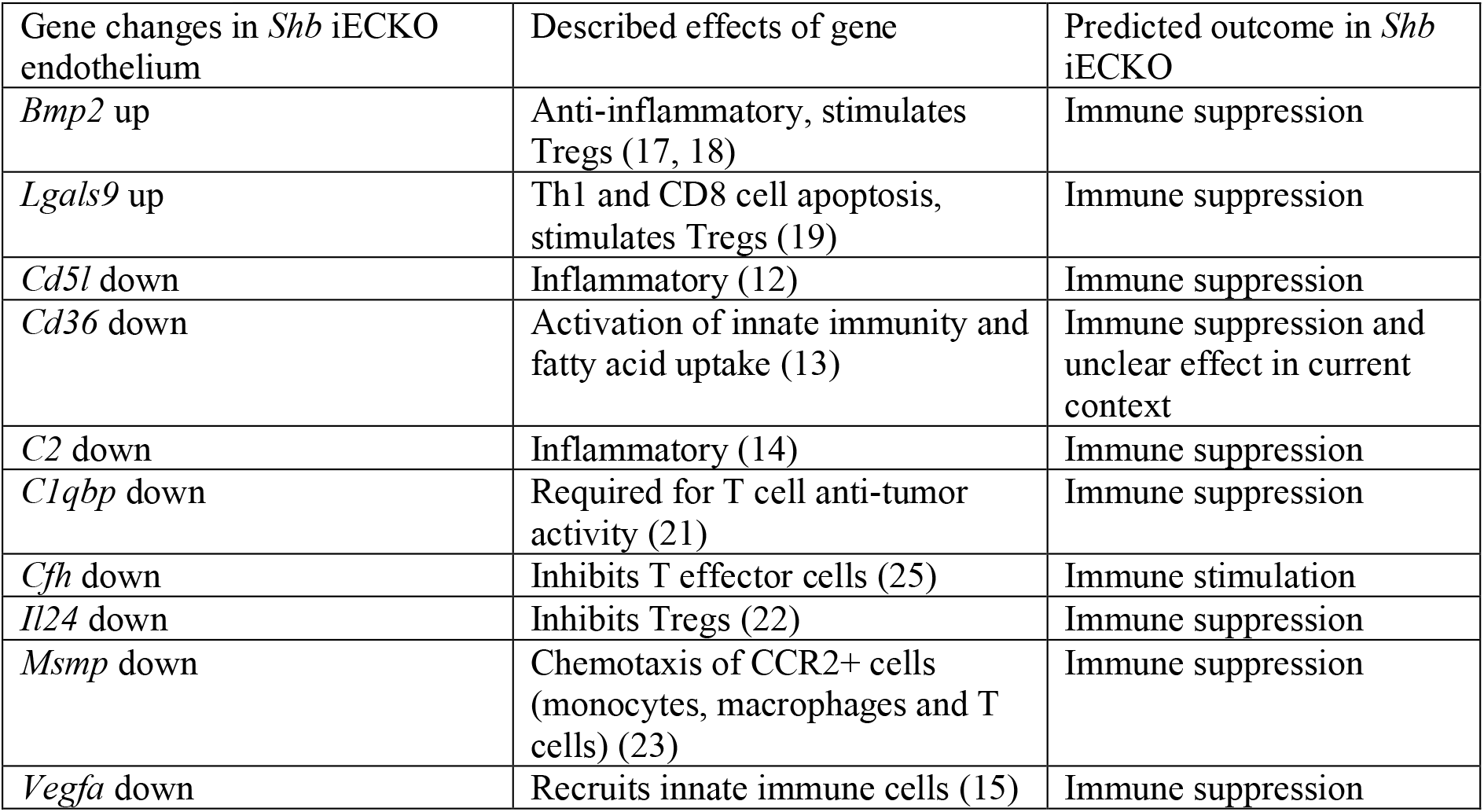
*Shb* iECKO gene expression changes curated from the categories “cytokine activity”, “positive regulation of T cell activation” and “activation of immune response” (see Table S1) pertaining to Gene Ontology biological processes of relevance for immune responses and T cell function not addressed in the previous publication (5).

We next addressed whether these gene expression changes are universal for vascular normalization or a specific consequence of *Shb*-gene inactivation. To address this issue, we compared *Shb* iECKO endothelium with normal breast endothelium (8) for the genes listed in Table 1 and those pertaining to TEM as previously published (5) in Table 2. Comparisons were made as described in the methods section. The human normal and breast cancer EC populations were arterial, capillary and venous. The rationale behind this approach was the assumption that normal endothelium largely reflects vascular normalization relative tumor endothelium and that the non-leaky, normalized vasculature due to *Shb* deficiency in EC represents an endothelial phenotype resembling that of healthy EC with both low leakiness and promotion of immune suppression. A potential agreement between the *Shb* iECKO relative human normal EC was noted for the combined arterial, capillary and venous endothelial gene expression profiles (Table 2, p<0.05 by Chi-square test comparing agreement with a random one-to-one response). Furthermore, there was a significant correlation between *Shb* iECKO relative normal arterial EC expression of these genes (Fig. 1A). However, no such correlation was observed when the corresponding ratios for all human EC populations were calculated (Fig. 1B). It should be noted that the relative number of venous EC analyzed in that study differed between the healthy and tumor groups, possibly confounding the results. However, *BMP2*, and *CD83* showed a dichotomy in their responses between the mouse and human sets and were thus classified as outliers (> 2 SD). When these two outliers were omitted, changes in *Shb* iECKO relative normal breast correlated (p<0.02) (Figure 1C) for the combined arterial, capillary and venous endothelial cell expression values. *CD83* exerts pleiotropic effects on the immune system depending on context (26) and consequently the discrepancy in response between the two comparisons may thus not necessarily imply a contradiction but rather reflect specific conditions that are distinct for the experimental mouse and human contexts. In addition, the *CD83* response showed an agreement between mouse EC normalization and normal human arterial (Table 2). *BMP2* is primarily anti-inflammatory but may also inhibit monocyte differentiation to type 2 (anti-inflammatory) macrophages (27). Gene expression differences between *Shb* iECKO relative normal human EC gene expression for genes well characterized as essential for leukocyte TEM (*SELE, SELP, VCAM1, ICAM1, ICAM2, F11R, JAM2, JAM3, CDH5, PECAM1, ESAM, CD99*) (28) were also presented. The values are based on average expression under the conditions irrespective of p-values. Regardless, there was a significant correlation between the two ratios (Fig. 1D), suggesting that the normalized *Shb* iECKO endothelium has a similar capacity for leukocyte TEM as the normal breast endothelium. Additionally, the differences in expression of these genes of importance for TEM between human control and breast cancer EC were limited (Table 3) among which *SELE, SELP* and *VCAM1* showed the most prominent changes and were approximately twice as high in normal venous EC as in tumor venous EC. This suggests that leukocyte TEM is not by itself a major component of an immune suppressive EC phenotype. Consequently, there was no difference in the number of CD45^+^-cells isolated from *Shb* iECKO and wild type tumors (5). In the human EC, only *ICAM2* and *CDH5* exhibited higher expression in arterial EC than venous EC (Table 3). Expression of the other genes of importance for TEM was similar or much lower in arterial EC, further supporting the well-established notion that the bulk of leukocyte TEM occurs at venular sites (28). To substantiate the concept of immune suppression further in normal arterial EC compared with breast cancer arterial EC, gene ontology of significantly altered genes related to cytokine activity showed highly significant changes in processes pertaining to stimulation of various immune-related processes in tumor EC (Table 4). Venous EC displayed a similar pattern of changes, albeit with higher p-values (results not shown). It can be noted that the genes increased in wild type tumor EC relative *Shb* iECKO in Table 2 were associated with the GO-term “positive regulation of immune system process” (p-value = 2.245E-8), further supporting the notion that *Shb* iECKO and normal breast EC confer similar immune suppressive features on the breast vasculature. In summary, the *Shb* deficient EC phenotype causing vascular normalization demonstrates an immune suppressive gene expression profile resembling that of normal breast, and particularly so, arterial EC.

**Table 2:**
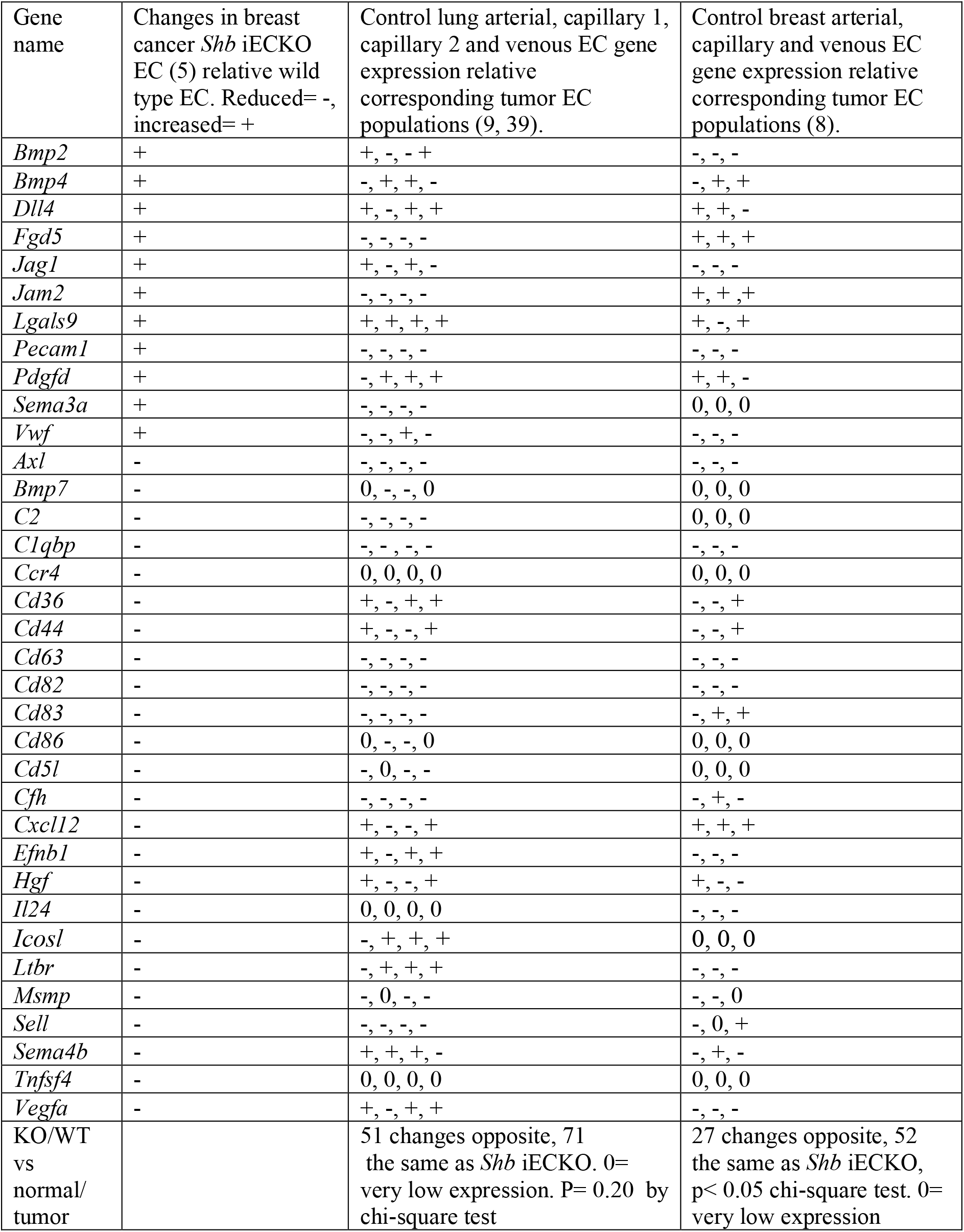
Comparison of *Shb* iECKO breast cancer endothelial gene expression changes with published control and tumor endothelial gene expression changes. Genes that exhibited low expression in the scRNAseq analysis (< 0.015 transcripts per million) were omitted as indicated by zero values and *CD5l* was not detected in these databases.

**Table 3:**
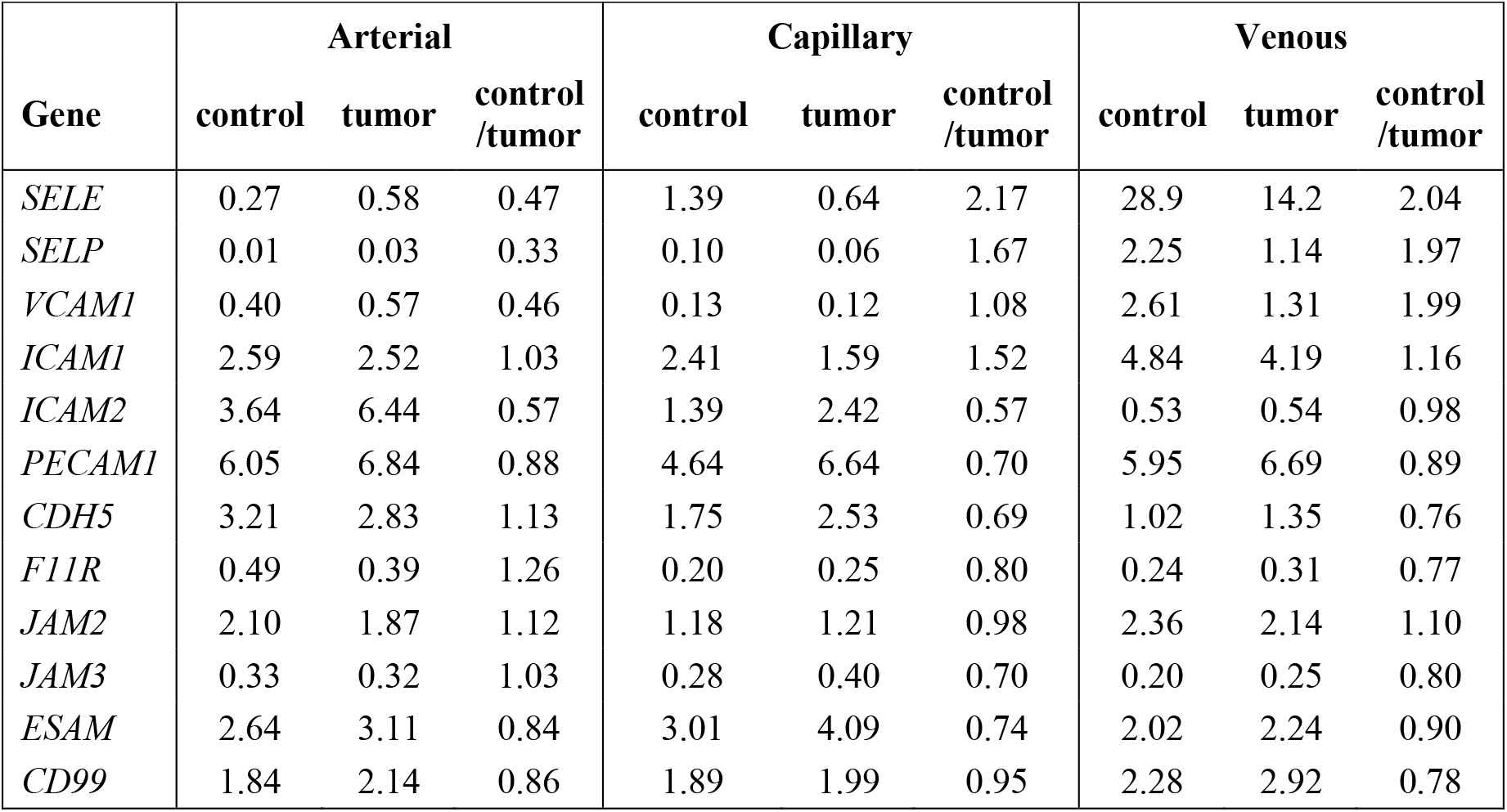
Human normal breast and breast cancer endothelial gene expression (in TPM) of selected genes of importance for leukocyte TEM in arterial, capillary and venous EC (8).

**Table 4:**
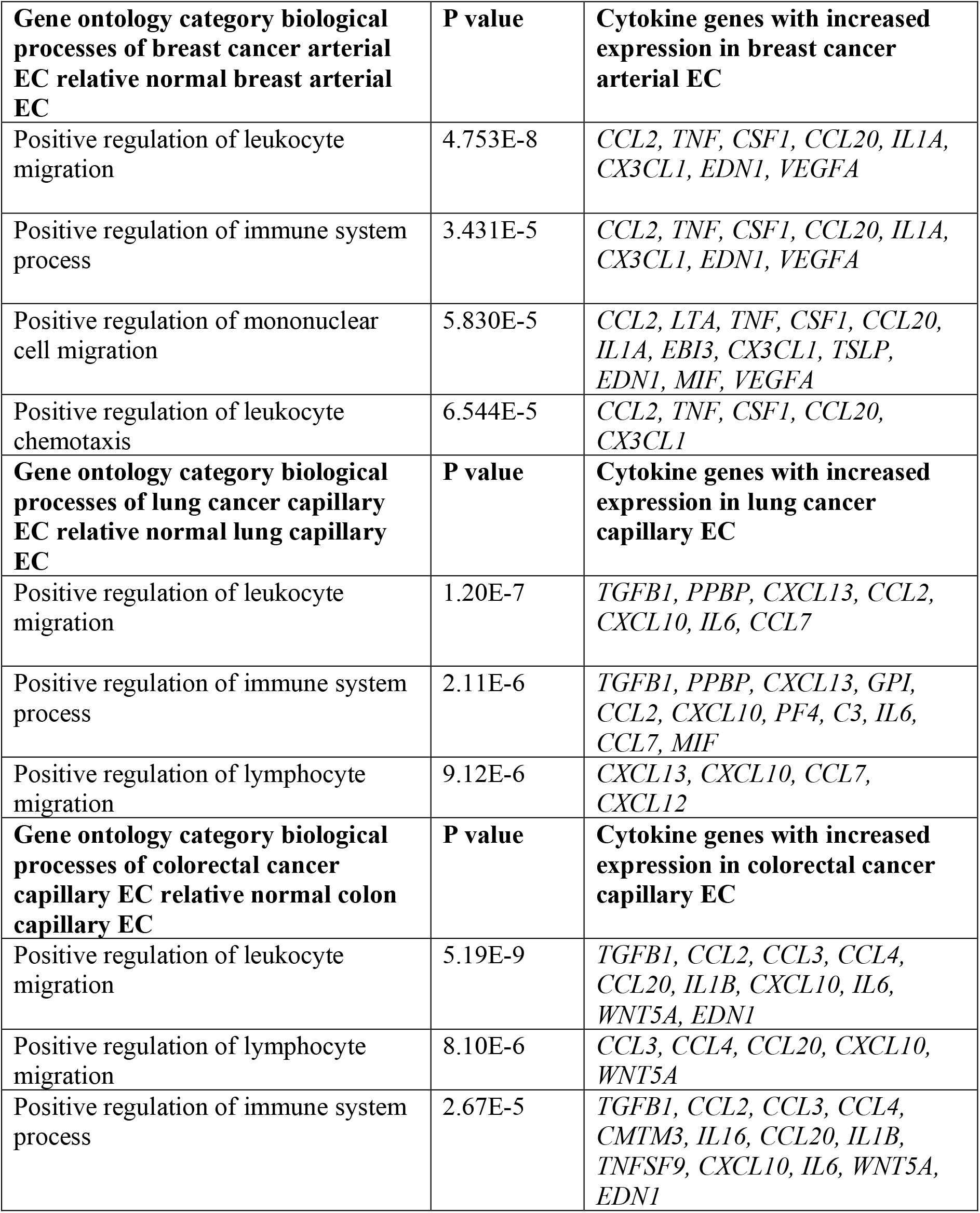
Gene ontology pertaining to immune responses when comparing breast cancer arterial, lung capillary (type I and II) EC or combined colorectal cancer EC populations with the corresponding normal EC populations for significantly (p< 0.05) altered genes related to cytokine activity after correction for multiple comparisons. The breast cancer venous EC population showed similar gene ontology changes as the arterial population but with higher p values.

**Figure 1.**
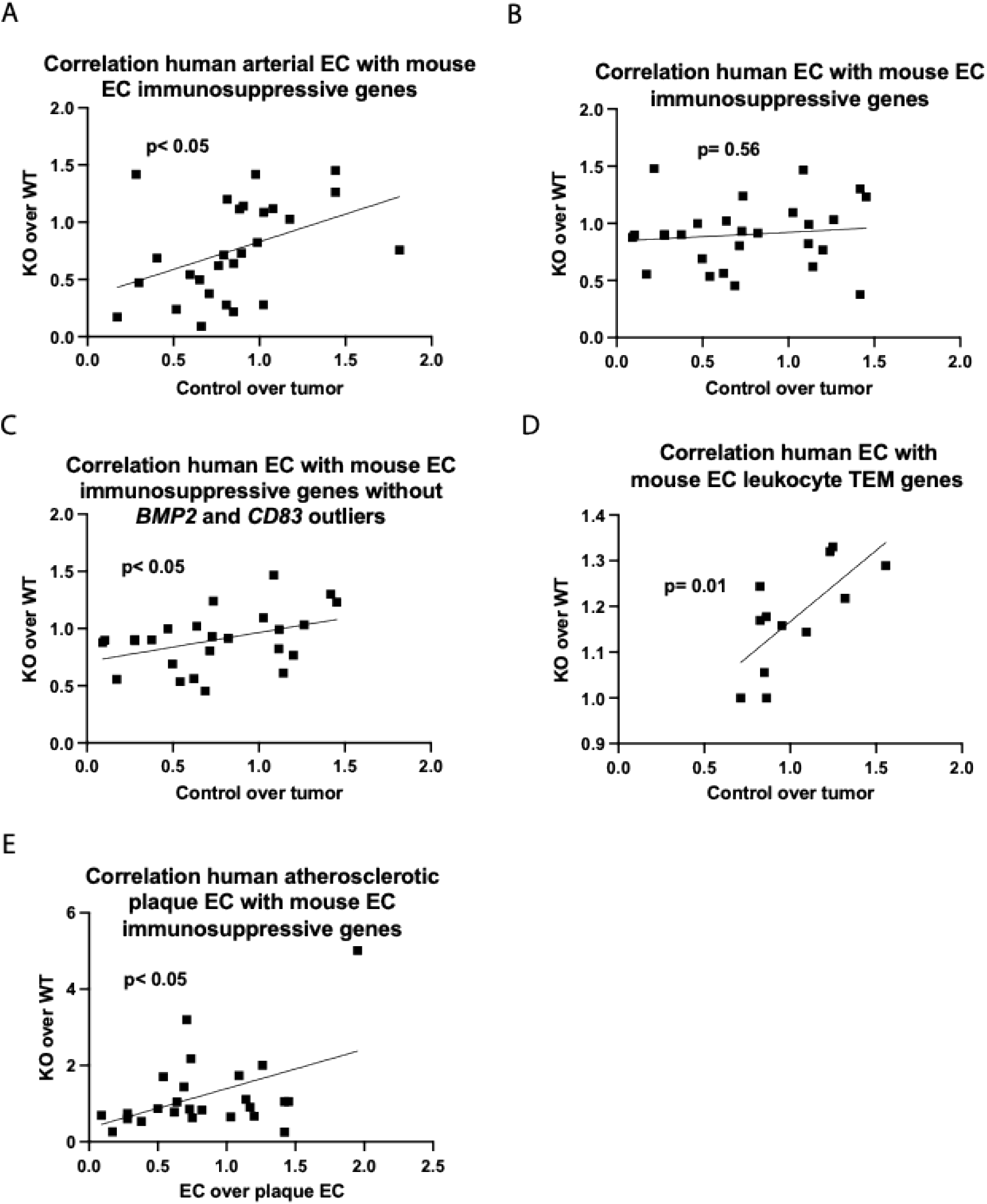
Correlation *Shb* iECKO over WT with control/tumor EC for the immune suppressive genes listed in Table 1. A) Human arterial EC compared with mouse EC gene changes. Genes with less than 0.015 TPM were excluded. B) Correlation all human EC compared with mouse EC gene changes. C) Correlation all human EC with mouse EC gene changes after omitting *BMP2* and *CD86* outliers. D) Correlation between *Shb* iECKO over WT with control/tumor EC for genes related to leukocyte TEM (*SELE, SELP, VCAM1, ICAM1, ICAM2, F11R, JAM2, JAM3, CDH5, PECAM1, ESAM, CD99*). E) Correlation normal EC over atherosclerotic plaque proangiogenic/endMT EC with *Shb* iECKO over WT EC immunomodulatory gene expression. Significantly altered genes were *AXL, BMP2, BMP4, C1QBP, C2, CD36, CD44, CD64, CD82, CFH, CXCL12, DLL4, EFNB1, FGD5, HGF, JAG1, JAM2, LTBR, MSMP, PDGFD, PECAM1, SELL, SEMA3A, VEGFA* and *VWF*.

As described above, the best correlation between mouse breast cancer normalized endothelium and human normal endothelium was observed for arterial EC. Consequently, we decided to investigate the proangiogenic and endMT EC gene expression profiles of human atherosclerotic plaques to address whether this correlation would be affected by pathological changes in arteries (11). Among the list of immune suppressive genes in Table 2, 25 were significantly altered in either of the proangiogenic and endMT sets, or both, EC populations in plaques. The normal EC gene expression for these genes was found to correlate significantly with the corresponding *Shb* iECKO EC gene expression changes (Fig 1E), suggesting a similarity in the immunomodulatory EC gene expression profile of atherosclerotic plaques and breast cancer.

Immunomodulatory gene expression changes are dependent on transcription factor alterations, and the study was thus extended by searching for relevant transcription factors potentially regulating the differences in the list of immune regulating genes (Table S2). A considerable overlap in transcription factor binding sites was noted for this set of genes and all but one (*CD36*) displayed potential binding sites for transcription factors that exhibited shared expression changes in *Shb* iECKO and human healthy EC. Search for enriched transcription factor binding to the immunomodulatory genes as listed in Table 2 revealed one transcription factors (*BCL6*) differentially expressed in both *Shb* iECKO and normal breast arterial EC that could contribute to 10 of the 35 changes in immunomodulatory gene expression.

The question remains whether the correlation between immune suppression and vascular normalization is specific for breast cancer or can be observed in other types of cancer. Comparison of lung cancer EC gene expression changes with those of normal lung endothelium showed that the agreement in gene expression changes between the *Shb* iECKO tumor endothelium relative normal lung endothelium was less apparent (Table 2). In this comparison, 71 changes were the same whereas 51 were opposite (p<0.20 when comparing with a random one-to-one response). However, since this is a different type of cancer, less agreement is expected. Gene ontology analysis, however, suggested that tumor-induced alterations in EC cytokine gene expression when compared with normal lung capillary (type I and II) EC conferred an immunostimulatory mode (Table 4) which is in line with the breast cancer data. The significant gene expression changes in arterial and venous EC were too few to perform gene ontology analysis. Similarly, the combined colorectal cancer EC populations displayed immunostimulatory changes (Table 4). Comparisons could only be made on the merged total EC population due to low power when individual EC populations were analyzed. It is concluded that tumorigenicity provokes immunostimulatory properties in EC in multiple cancers.

## Discussion

The data suggest that vascular normalization confers an immune suppressive phenotype in breast cancer. The notion that normalized tumor endothelium in breast cancer supports immune suppression may appear counterintuitive and it has been reported that untreated tumor vasculature is immunosuppressive (29). Consequently, combining treatment causing vascular normalization with immunotherapy commonly enhances the response to immunotherapy (30). An alternative interpretation to that finding is that when the tumor vasculature becomes normalized and immune suppressive, immunotherapy shows increased efficacy. The relative increase in Tregs in breast tumors with normalized vasculature previously reported (5) would thus make the tumors more amenable to immune checkpoint inhibition. From a physiological perspective, homeostatic mechanisms may be operating under normal conditions that prevent autoimmunity and the immune suppressive features of normal endothelium could belong to that category. Upon infection, inflammation ensues allowing leakage and immune cell extravasation and this could result in a lower population of extravasated Tregs, thus promoting an activation of the immune response against the pathogen. In many respects, the tumor vasculature resembles an inflamed vasculature (31) and thus one can consider similar mechanisms operating in a tumor microenvironment as in infection. However, tumor EC have been reported to be immunosuppressive since they may exhibit increased expression of the immune checkpoint protein gene *CD274* (32) and show decreased expression of the cell surface adhesion molecule *ICAM1* that has been shown to increase upon VEGFA-targeting therapy (33). On the other hand, EC activated by inflammation support immune responses by producing inflammatory cytokines/chemokines and increase their expression of cell surface adhesion molecules allowing increased leukocyte TEM (34). This dichotomy is difficult to explain but it can be noted that VEGFA-targeting may cause tumor necrosis (33) and thus such features need to be taken into consideration to obtain a complete understanding of the response in a tumor context. In the breast cancer study currently analyzed (8) there was no evidence for increased expression of *CD274* (ratio normal over breast cancer EC 1.19) or altered expression of *ICAM1* (Table 2) in the tumor venous endothelium arguing against the notion that breast cancer EC are immunosuppressive by themselves. Consequently, a high leakage response, which is a common feature in tumors, would thus be immune supportive and prevent early tumor expansion in many cases. As the tumor evolves, tumor intrinsic factors are likely to promote immune suppression unrelated to and counteracting the immunostimulatory propensity of an otherwise inflamed and leaky endothelium. Instances in which tumor EC display immune suppressive properties could reflect the release of factors from malignant cells that operate in a context of promoting an immunosuppressive microenvironment. Due to tumor heterogeneity, one might expect a plethora of mechanisms exhibiting immunosuppression with respect to both EC and immune cell functions under such conditions. It should be noted that VEGFA may also exert direct immune suppressive effects on dendritic and T cells (16) and this will contribute to the final balance between immune suppressive and immune stimulatory cues in the tumor microenvironment independently of EC.

The healthy Arteria Intima and Media are considered immune privileged due to combined effects of EC and vascular smooth muscle cells (35) and endothelial dysfunction contributes to plaque formation (36). Therefore, it is not surprising that the *Shb* iECKO gene expression profile correlated best with normal breast arterial EC immune regulatory gene expression, assuming that *Shb* iECKO EC indeed are immune suppressive. The correlation pattern of arterial EC immune suppressive gene changes with the *Shb* iECKO profile suggests that these do not directly pertain to TEM but are related to separate immune suppressive features of the endothelium. However, atherosclerosis is a disease in which blood monocytes adhere to and cross the endothelial barrier at sites with shear stress-exposed endothelium in susceptible arteries, thus initiating an inflammatory response in the subendothelial Tunica Intima (37). This insult will eventually involve a T cell response with a gradual decrease of Tregs and increase of Th2 cells that aggravate disease (38). Intuitively, one can consider an immune suppressive gene expression profile in arterial endothelial cells as advantageous in preventing lesion formation. The data in Fig 1F support this notion.

In conclusion, vascular normalization in a mouse experimental model of breast cancer confers an immune suppressive gene expression profile that resembles that of normal human breast endothelium. A possible interpretation would be that normal endothelium in many tissues is immune suppressive and that inflammation caused by tumors, pathogens and other sources switches the endothelial phenotype to an immune stimulatory mode. Tumors develop additional strategies that bypass the normal immunoreactive features of an inflammatory vasculature to overcome immunity. This concept presents a therapeutic conundrum. Although a low-leaky tumor vasculature is desirable from a therapeutic perspective, an immunostimulatory mode is also preferred. To find regimens that combine immune checkpoint inhibition with measures that simultaneously promote EC inhibition of leakage and immune stimulation is a task for future studies.

## Abbreviations

EC: endothelial cells
TEM: transendothelial migration
SC: single cell
VEGFA: vascular endothelial growth factor -A
GO: gene ontology
SHB: Src-homology domain protein B
iECKO: induced conditional deletion in EC by tamoxifen injections using *Cdh5-CreERT2* transgene
endMT: endothelial-mesenchymal transition
Treg: T regulatory cells
Th2: T helper 2 cell
TF: transcription factor.

## Acknowledgements

We are grateful to Dr Lena Claesson-Welsh for comments. The computations/data handling was enabled by resources in project UPPMAX 2025/2-491 provided by Uppsala University at UPPMAX.

## Legends to supplementary tables

Table S1: Immune regulatory genes with p< 0.05 in *Shb* iECKO endothelium not previously identified (5). Genes responsible for positive regulation of T cell activation, immune responses, cytokine activity were curated according to (https://www.informatics.jax.org/vocab/gene_ontology). Only gene products secreted or expressed on endothelial cell surface were included. Red; increased expression, blue; decreased expression.

Table S2: Transcription factor alteration between normal human breast and human breast cancer endothelial cells. In “shared transcription factors”, the target genes are listed. Transcription factors (TF) altered and increased for human arterial, capillary, venous and *Shb* iECKO are listed as well as those shared between the gene sets. In “TF gene target binding sites”, potential binding of the set of transcription factors to the target genes are listed. “Enriched TF for the target genes” shows transcription factors with enriched binding potential among the target genes. *BCL6* stands out as a transcription factor that displayed significantly different expression between the conditions with enriched binding to 10 of the target genes.

